# Versatile polyketide biosynthesis platform for production of aromatic compounds in yeast

**DOI:** 10.1101/618751

**Authors:** Tadas Jakočiūnas, Andreas K. Klitgaard, David Romero-Suarez, Christopher J. Petzold, Jennifer W. Gin, Yaojun Tong, Rasmus J.N. Frandsen, Tilmann Weber, Sang Yup Lee, Michael K. Jensen, Jay D. Keasling

## Abstract

To accelerate the shift to bio-based production and overcome complicated functional implementation of natural and artificial biosynthetic pathways to industry relevant organisms, development of new, versatile, bio-based production platforms is required. Here we present a novel yeast platform for biosynthesis of bacterial aromatic polyketides. The platform is based on a synthetic polyketide synthase system enabling a first demonstration of bacterial aromatic polyketide biosynthesis in a eukaryotic host.

## Introduction

An increasing number of chemicals are being produced by environmentally-friendly bio-based synthesis^1,2^ to overcome the problems of low-yielding chemical synthesis or solvent-heavy extraction from natural resources, for achieving a sustainable way of life. Unfortunately, development of microbial cell factories for the bio-based production of desired chemicals often requires significant amount of time, resources and efforts to meet industrial demand, hence the shift towards bio-based production is slow. Also in many cases native hosts are not suitable for industrial conditions due to low production level and/or complicated culturing conditions, necessitating the use of a heterologous hosts such as *Escherichia coli* or yeast^3,4^. However, production of natural products, such as polyketides, in heterologous hosts has proven difficult or even impossible^5–7^. To overcome these limitations, standardized, versatile and programmable biosynthesis platforms in genetically traceable, and robust hosts are desired.

Polyketides are a large class of bioactive natural compounds found widely in fungi (type I iterative and type II), bacteria (mostly type II, but also type I modular and type III) and plants (type III), possessing a variety of biological activities, including antibacterial, anticancer, antifungal, antiviral and many more^8–11^. As a consequence, polyketides have been, and still are, major leads in drug discovery programs^11–14^.

The most diverse and widely studied polyketides are originating from bacteria^11^. Complex bacterial aromatic polyketides can be produced through non-reducing polyketide pathways, where two-carbon units (-CH2-CO-, ketides) are polymerized into linear polyketide chains of various length by multicomponent enzyme complexes known as polyketide synthase (minimal PKS), and then can be folded to form aromatic structures^15–17^. Folding in most bacterial systems is facilitated by aromatases and cyclases, and the resulting products are further modified by other classes of tailoring enzymes that closely interact with the minimal PKS^18^. From the bioengineering point of view, bacterial type II PKS systems offer flexibility in terms of choice from vast amount of aromatases, cyclases and tailoring enzymes to allow for the rational engineering of pathways to form desirable aromatic compounds^19^ and to develop programmable polyketide production platforms. Development and optimization of such a production platform in native bacterial hosts can be troublesome due to the lack of genetic tools, production of unwanted toxic metabolites, complicated culturing conditions, and low or conditional production of desired compounds^3,4,20^. Unfortunately, bacterial type II PKSs have not yet proven possible to express in eukaryotes^15^. In contrast, plant (type III) PKSs, that ultimately form aromatic compounds, via linear non-reduced polyketide intermediates, consist of a single enzyme^21^ and can be expressed in heterologous eukaryotic hosts to form a polyketide chain of varied length, yet the lack of characterized cyclases, aromatases and tailoring enzymes in plant PKS systems limits the use of type III PKSs for versatile polyketide biosynthesis^22,23^. Although, recently it was demonstrated that it is possible to functionally combine the activity of plant type III PKS with bacterial type II PKS related cyclase and aromatase in plants and filamentous fungi^23,24^

Here, we describe a first-of-its-kind programmable polyketide production platform in the yeast *Saccharomyces cerevisiae*, based on combining the synthesis of a polyketide (octaketide) by plant-based type III octaketide synthase (OKS) from *Aloe arborescens* to produce type II polyketide products from benzoisochromanequinone antibiotic – actinorhodin pathway (Act) from *Streptomyces coelicolor*.

## Results

### Expression of Act gene cluster in yeast

First, we reconstructed a widely studied actinorhodin pathway^25^ by integrating required codon optimized genes into the yeast genome together with its bacterial minimal PKS (Figure 1A). Gene assemblies and genomic integrations were performed in 2-3 steps by first performing *in vivo* assembly of expression units in *Escherichia coli*, and second by using our recently developed CRISPR/Cas9 genome engineering techniques to integrate the assembled gene expression units into the yeast genome (Supplementary Table 1)^26,27^. Since actinorhodin and other intermediates in the pathway have color^28^, successful production through the pathway was initially expected to be assessed by visual inspection of engineered yeast. However, from the first designs, no apparent or very modest color was observed in yeast cells harboring actinorhodin pathway (Supplementary Figure 1; TC-140, TC-156). To mitigate the lack of (or modest) visual phenotypes we next investigated if all proteins from the Act pathway were successfully expressed using whole cell proteomics. From this analysis it was evident that most of Act proteins were detected except for ActVI-2 (dehydrogenase) and ActI-1 (3-oxoacyl-ACP synthase), the latter of which is needed for the first committed step of the minimal ActPKS (Supplementary Figure 2). Further, we aimed to elucidate which metabolites, if any, are produced from first generation Act pathway design. Since none of the reported pathway metabolites are commercially available as standards, we performed comparative LC-MS metabolite profiling using wild type *S. coelicolor* whole cell extract as a standard. This analysis indicated that none of the described intermediates from the Act pathway or Act itself were detected in the engineered yeast strains (Figure 1B; TC-140, TC-156), hinting that the Act – type II PKS indeed was not functionally expressed or correctly assembled into a functional PKS in yeast.

**Figure 1.**
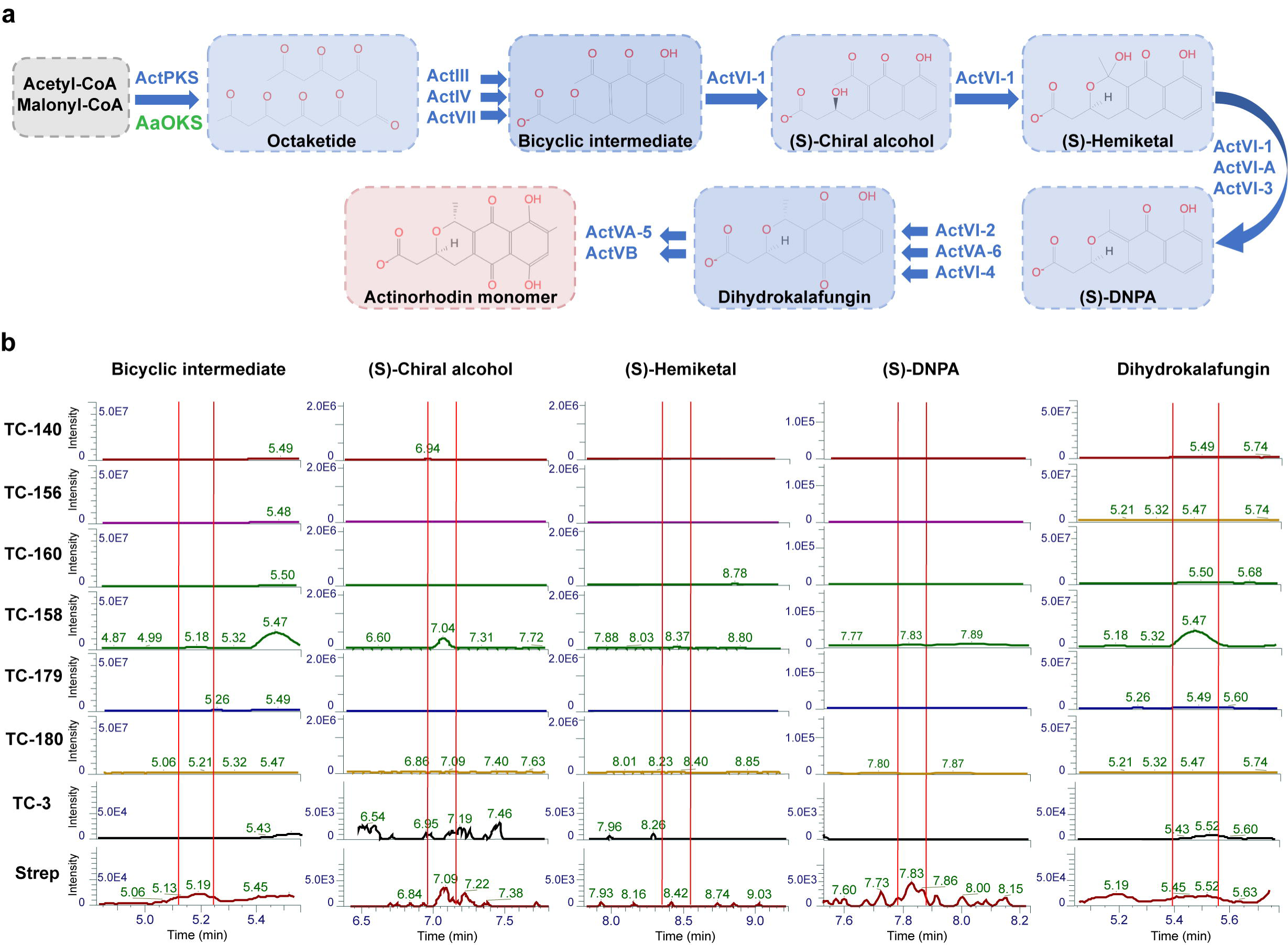
**a**, Schematic overview of actinorhodin pathway with Act minimal PKS or with *Aa*OKS expressed in yeast, including enzymes and chemical compounds produced through the pathway. Enzymes are listed in blue and green, blue arrows depict chemical reactions catalyzed by listed enzymes and produced compounds are depicted in blue or red shade. **b**, Chromatograms from comparative LC-MS metabolomics showing investigated metabolites in yeast. The main products or intermediates investigated by LC-MS were bicyclic intermediate, (S)-chiral alcohol, (S)-hemiketal, (S)-DNPA, dihydrokalafungin (DHK). Metabolites from the natural actinorhodin producer *S. coelicolor* were used as a standard (positive control) for comparing metabolites produced in *wt* and engineered yeast strains: TC-3, TC-140, TC-160, TC-156, TC-158, TC-179, TC-180. Red vertical lines on chromatograms depict the peaks of listed compounds based on known mass and positive control. Intensities of the peaks and elution times are shown on the corresponding axis.

**Figure 2.**
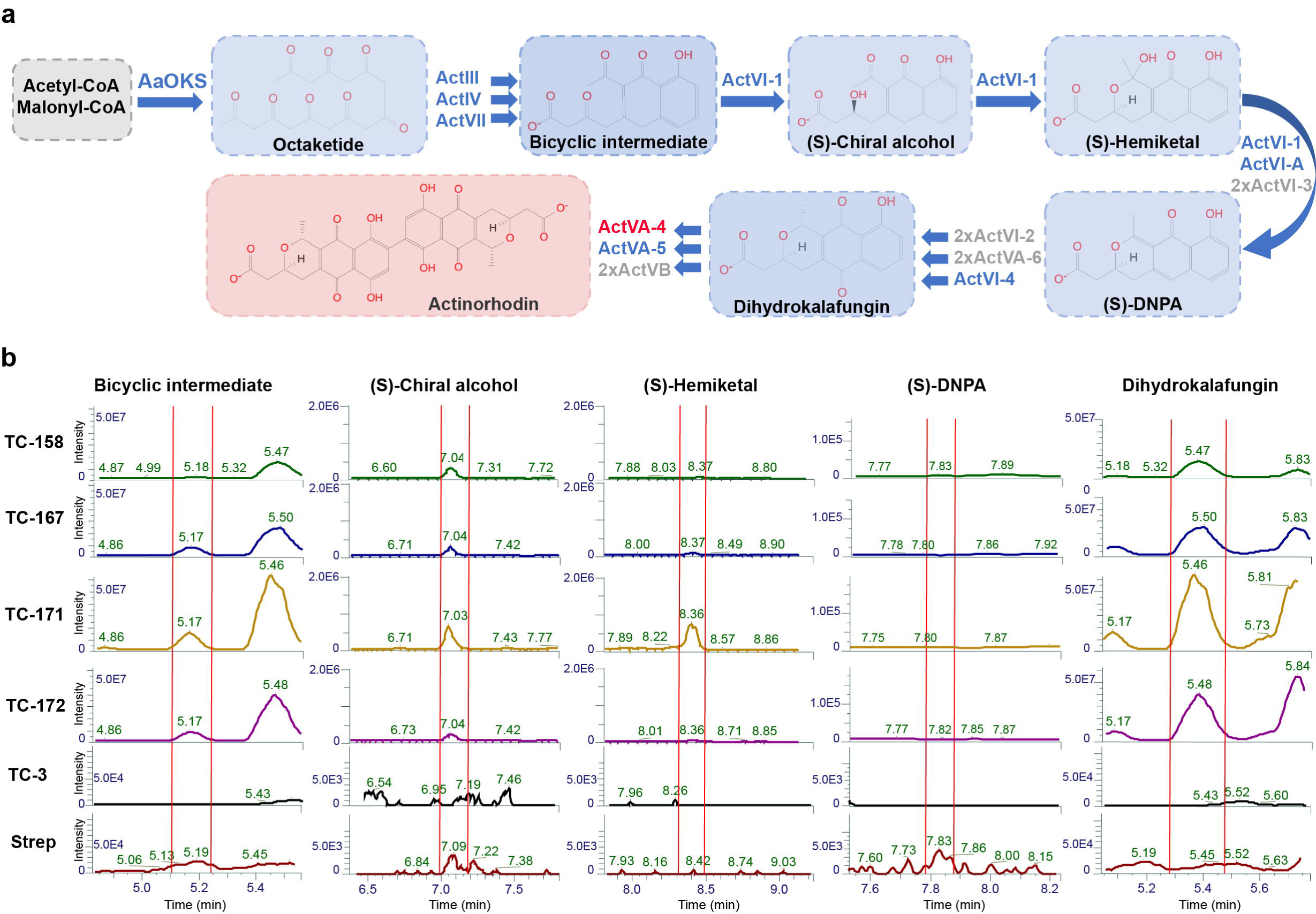
**a**, Schematic overview of the optimized actinorhodin pathway where the Act minimal PKS is replaced with *Aa*OKS in yeast, including enzymes and chemical compounds produced through the pathway. Enzymes encoded by a single-copy gene are indicated in blue, and enzymes encoded by multi-copy genes are indicated in grey. Blue arrows depict chemical reactions catalyzed by listed enzymes and produced compounds are depicted in blue or red shade. Dimerize required for dimerization of DHK to form actinorhodin is deciphered in red. **b**, Chromatograms from comparative LC-MS metabolomics showing investigated metabolites in optimized yeast strains. Main products or intermediates were investigated by LC-MS: bicyclic intermediate, (S)-chiral alcohol, (S)-hemiketal, (S)-DNPA, dihydrokalafungin (DHK). Metabolites from natural actinorhodin producer *S. coelicolor* (deciphered as Strep) was used as a standard (positive control) for comparing metabolites produced in *wt* and engineered yeast strains: TC-3, TC-158, TC-167, TC-171, TC-172. Red vertical lines on chromatograms depict the peaks of listed compounds based on known mass and positive control. Intensities of the peaks and elution times are shown on the corresponding axis.

### Replacement of Act minimal PKS with *Aa*OKS

To overcome the lack of function of type II PKS we replaced the Act minimal PKS with a type III octaketide synthase from plant *A. arborescens* (*Aa*OKS)^29^, which was described to produce a polyketide product with an identical chain-length as Act. Hence, the second generation production strains were created by replacing the actinorhodin minimal PKS with *AaOKS*, but retaining the rest of the actinorhodin pathway (Figure 1a). The first strain design expressing *AaOKS* indeed displayed pigmented colonies indicating the production of Act cluster metabolites (Supplementary Figure 1; TC-158). We further investigated expression optimization of *AaOKS* by integrating either a single-copy of the *AaOKS* gene into the genome or expressing the *AaOKS* from a high-copy plasmid. As judged from phenotypic inspection, only the Act pathway strain expressing *AaOKS* from a high-copy plasmid gave rise to pigmented colonies potentially derived from Act pathway metabolites (Supplementary Figure 1; TC-160 vs. TC-158). Further, from comparative metabolite profiling, the four main products (bicyclic intermediate, (S)-chiral alcohol, (S)-hemiketal, dihydrokalafungin) from the Act pathway were tentatively observed in this strain (Figure 1b; TC-158), while no production was observed in the *wt* control (TC-3), or strains i) expressing Act pathway without a PKS (TC-156), ii) expressing Act pathway and single-copy of *AaOKS* (TC-160), or *wt* control expressing *AaOKS* in either single- or high-copy (TC-179 or TC-180) without the Act pathway (Figure 1b). The other major intermediate (S)-DNPA from the Act pathway could not be detected or confirmed reliably, most likely because this product was metabolized rapidly by Act pathway enzymes. By comparative LC-MS analyses, based on known mass and published UV data^30^, and direct comparison to *S. coelicolor* metabolites, we observed accumulation of a compound expected to be dihydrokalafungin (DHK), as a final product (Figure 1b; TC-158). DHK is known to possess anticancer, antifungal, antiparasitic properties, and it is a potential drug target^31,32^. To further investigate the compound putatively identified as DHK, we performed more thorough analysis including LC-MS/MS, as no standards were commercially available. These results indicated that DHK was indeed being produced in the engineered yeast strain as its MS/MS fragmentation pattern was the same in both engineered yeast strain and *S. coelicolor* (Supplementary Figure 3).

**Figure 3.**
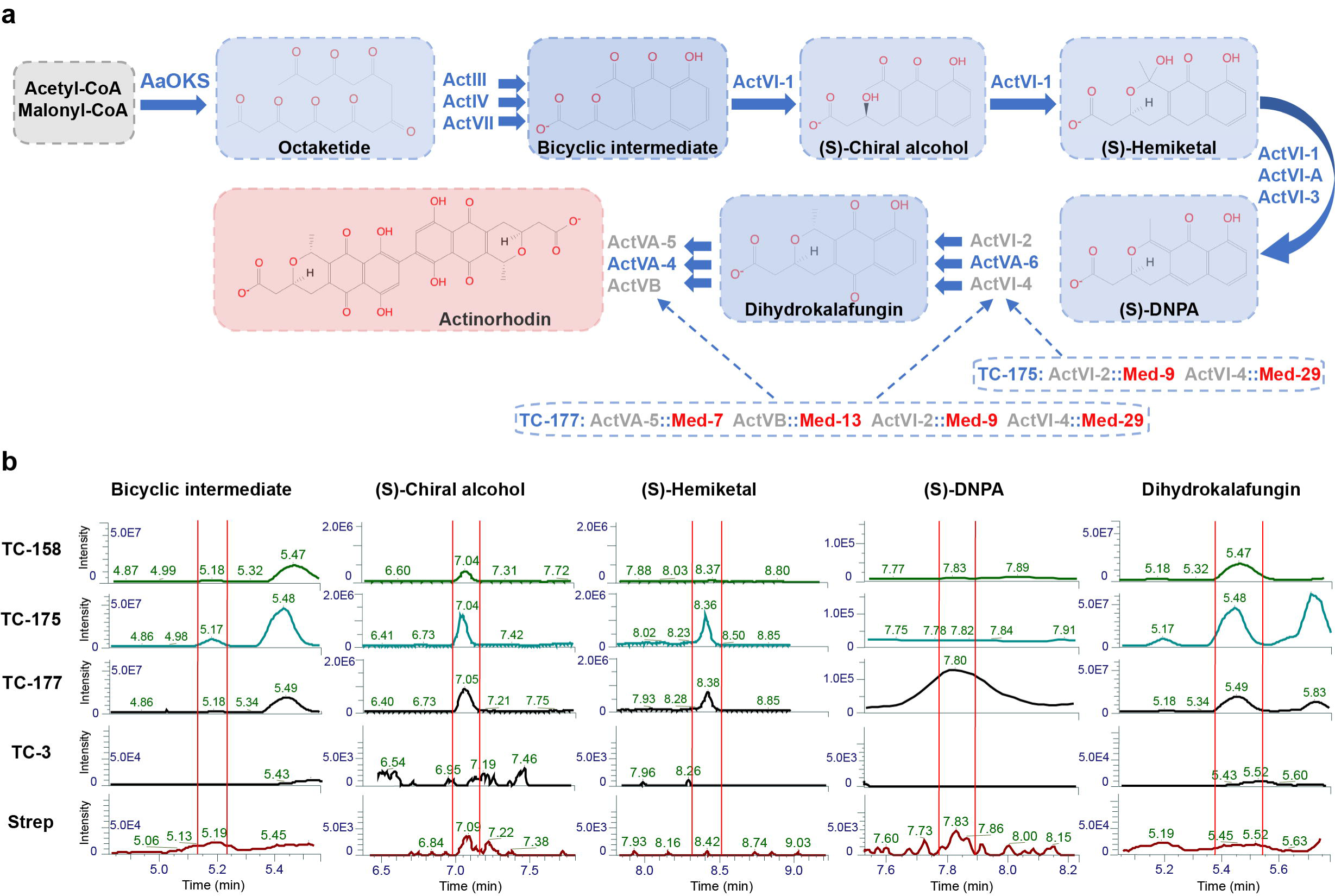
**a**, Schematic overview of the re-programmed actinorhodin pathway where ActVI-2, ActVI-4, ActVA-5, ActVB (native Act pathway enzymes) were functionally replaced with enzymes Med-9, Med-29, Med-7, Med-13 respectively (from medermycin biosynthetic pathway) to test the versatility and aprogramability of the heterologous pathway. Enzymes are listed in blue and Act pathway enzymes which were replaced in grey; enzymes which were newly integrated in to yeast genome shown in red. Blue arrows depict chemical reactions catalyzed by listed enzymes and produced compounds are depicted in blue or red shade. **b**, Chromatograms from comparative LC-MS metabolomics showing investigated metabolites in reprogrammed yeast strains. Main products or intermediates were investigated by LC-MS: bicyclic intermediate, (S)-chiral alcohol, (S)-hemiketal, (S)-DNPA, dihydrokalafungin (DHK). Metabolites from the natural actinorhodin producer *S. coelicolor* (indicated as Strep) was used as a standard (positive control) for comparing metabolites produced in *wt* and engineered yeast strains: TC-3, TC-158, TC-175, TC-177. Red vertical lines on chromatograms depict the peaks of listed compounds based on known mass and positive control. Intensities of the peaks and elution times are shown on the corresponding axis.

### Optimization of aromatic polyketide production platform

To optimize the platform strain further for production of type II polyketide compounds, we next integrated a second copy of each of the four genes encoding ActVI-3, ActVI-2, ActVA-6, ActVB (Figure 2a), all showing low abundances as evaluated from whole cell proteomics analysis (Supplementary Figure 4). Upon overexpression of the four genes, the colonies became more intensely colored (Supplementary Figure 1; TC-167). To investigate if this phenotype could be correlated with increased DHK production, we performed comparative metabolite profiling by LC-MS, and noted that production of the DHK was indeed increased (Figure 2b; TC-158 vs. TC-167).

Next, we aimed to produce one of the most widely studied model antibiotics, actinorhodin, (a dimer of DHK) in both non-optimized (TC-158) and optimized (TC-167) yeast platform strains. DHK dimerization, as previously described^30^, is potentially catalyzed by the enzyme ActVA-4 (Figure 2a), which initially was not introduced into the yeast platform strains. Introduction of the dimeraze should allow for dimerization of DHK and production of the actinorhodin. However, as judged by the color of yeast colonies (Supplementary Figure 1; TC-171 with the non-optimized Act pathway and TC-172 with the optimized Act pathway), no significant changes were observed after introduction of the dimerase. Further analysis by comparative LC-MS corroborated the phenotypic result revealing no detectable actinorhodin in the yeast strains (Supplementary Figure 5). In addition, actinorhodin and its intermediates can also be toxic to yeast, and accumulated amounts could inhibit cell growth, and potentially compromise actinorhodin production. We tested this hypothesis by growing *wt* yeast in serial dilutions of conditioned medium where *S. coelicolor* had previously been grown. According to our LC-MS analysis (Figure 1b), this medium contained Act pathway metabolites together with actinorhodin. Here it was observed that growth of yeast cells was completely inhibited in 2x and 4x diluted conditioned medium, and even modestly compromised in 8x diluted conditioned medium (Supplementary Figure 6), while yeast cultivated in standard ISP2 medium displayed normal growth behaviour, thus indicating toxicity of Act pathway metabolites. Further, it was also observed that engineered yeast strains, producing DHK, showed reduced growth, as judged by colony size and growth profiling (Supplementary Figure 1; strains: TC-158, TC-167, TC-171; Supplementary Figure 7).

### Versatility and programmability of the system

The main goal of this study was to create a versatile and programmable platform for production of bacterial aromatic polyketides. To prove that our polyketide production platform can be engineered to express different polyketide synthesis modules, we replaced several key Act enzymes with enzymes from other *Streptomyces* species to achieve production of desired products (Figure 3a). For this purpose, we reconstructed the Act pathway in yeast so that ActVI-2 (dehydrogenase), ActVI-4 (dehydrogenase), ActVA-5 (hydroxylase) and ActVB (flavin□:□NADH oxidoreductase) from the Act pathway were replaced with enzymes Med-9 (dehydrogenase), Med-29 (dehydrogenase), Med-7 (oxygenase) and Med-13 (oxidoreductase), respectively, from the medermycin biosynthesis pathway (Med pathway)^33^ (Figure 3a). Next, we first phenotypically assessed the reprogrammed yeast strains (Supplementary figure 1; TC-175, TC-177), which provided an indication if the enzymes can be functionally replaced in the platform strains. Further, we investigated production of desired compounds with comparative metabolite profiling by LC-MS. LC-MS analysis revealed that detected metabolites are indeed the Med pathway intermediates in both strains with 2-4 enzymatic steps replaced by enzymes from Med pathway (Figure 3b; TC-175, TC-177). These results indicate that the developed platform system can be reprogrammed and employed for production of diverse bacterial polyketide compounds.

## Discussion

In summary, we have developed a functional first-of-its-kind eukaryotic production platform for bacterial polyketide derived products by employing a plant type III polyketide synthase to produce compounds originally found only in bacteria. As a proof-of-concept we engineered and optimized our platform in *S. cerevisiae* to successfully produce a compound identified as bioactive bacterial polyketide DHK. Finally, we further demonstrated versatility and programmability of our system by replacing key enzymes in Act pathway with enzymes from different *Streptomyces* species to produce desired products. Such characterisation confirms ability of the system to functionalize and produce octaketide derived products in a well-described, eukaryotic production work-horse. We envision our platform to be useful for production of many novel compounds, including novel antibiotics, characterization and functionalization of them, and moreover, sustainable production through cell factories.

## Methods

### Strains, plasmids and media

The yeast strains used here were isogenic to CEN.PK2-1C. Strains and plasmids are listed in Supplementary Tables 2 and 3, respectively. Yeast cells were grown in complete medium (YPD) with 2% glucose and synthetic complete (SC) from Sigma, supplemented with 2% glucose. *E. coli* strains were propagated in LB medium supplemented with 200 mg of ampicillin, Streptomyces strain was grown in ISP2 medium with 4% glucose.

All primer names and sequences are listed in Supplementary Table 4.

### Plasmid and strain construction

To create polyketide expressing strains large set of genes were integrated by using advanced CRISPR/Cas9 technology^26,27^. To create expression cassettes, respective genes were codon optimized for yeast and ordered (Integrated DNA Technologies) as gene blocks. Gene blocks were amplified using corresponding primers (Supplementary Table 4) and first USER cloned with single or bi-directional promoters to yeast integrative plasmids as described in previously published method^27^. All created integrative plasmids with corresponding expression units are listed in Supplementary Table 3. By employing previously detailed procedure^27^, all integrative plasmids were linearized and with their corresponding gRNA plasmids transformed to yeast expressing Cas9 for integration of desired genes to the genome. Due to large number of genes to be integrated, this has been processed in 2-3 steps. In a single transformation 4-6 genes were introduced and created strain used for the next round of transformation until all pathway genes were integrated. All the other plasmids and strains were created in the same way as previously described^27^.

### Metabolite extraction

Yeast cells were cultured in 50 ml SC selective or YPD medium for 168 hours at 30°C, shaking 250rpm. Streptomyces cells were grown in ISP2 medium for 168 hours at 30°C, shaking 250rpm. Cells were then collected by centrifugation and resuspended in ethyl acetate, which was acidified with 10% acetic acid. Resuspended cells where mixed with 0,5 µM acid washed glass beads (Sigma) and bead beated to break the cells. Lysates were centrifuged and upper ethyl acetate layer with extracted metabolites was collected. Ethyl acetate was evaporated and extracts resuspended in methanol for further analysis by LC-MS.

### Whole cell proteomics

The yeast cultures were grown in YPD medium in triplicates. Exponentially growing cells were harvested (totally OD_600_ – 20) and cell pellets flash-frozen in liquid nitrogen.

To prepare for protein lysis and precipitation, the yeast cell pellets were treated with 0.5 uL (2.5 U) of Zymolyase in 200 uL of 1 M Sorbitol 0.1 M EDTA at 37 °C for 30 min to digest cell walls and centrifuged at 20,817 × g for 1 minute. The supernatant was removed before continuing with a chloroform-methanol extraction as described previously, which was achieved by the addition of 80 µL of methanol, 20 µL of chloroform, and 60 µL of water, with vortexing followed by centrifugation at 20,817 × g for 1 minute to induce phase separation. The methanol and water layer was removed and then 100 µL of methanol was added and the sample with vortexing briefly followed by centrifugation for 1 minute. The chloroform and methanol mixture was removed by pipetting to isolate the protein pellet. The protein pellet was resuspended in 100 mM ammonium bicarbonate (AMBIC) with 20% methanol and quantified by the Lowry method (Bio-Rad DC assay). A total of 100 µg of protein was reduced by adding tris(2-carboxyethyl)phosphine (TCEP) to a final concentration of 5 mM for 30 minutes, followed by alkylation by adding iodoacetamide at a final concentration of 10 mM with incubation for 30 mins, and subsequently digested overnight at 37 °C with trypsin at a ratio of 1:50 (w/w) trypsin:total protein.

Peptides were analyzed using an Agilent 1290 liquid chromatography system coupled to an Agilent 6460QQQ mass spectrometer (Agilent Technologies, Santa Clara, CA) operating in MRM mode. Peptide samples (10 µg) were separated on an Ascentis Express Peptide ES-C18 column (2.7 µm particle size, 160 Å pore size, 50 mm length × 2.1 mm i.d., 60 °C; Sigma-Aldrich, St. Louis, MO) by using a chromatographic gradient (400 µL/min flow rate) with an initial condition of 95% Buffer A (99.9% water, 0.1% formic acid) and 5% Buffer B (99.9% acetonitrile, 0.1% formic acid) then increasing linearly to 65% Buffer A/35% Buffer B over 5.5 minutes. Buffer B was then increased to 80% over 0.3 minutes and held at 80% for two minutes followed by ramping back down to 5% Buffer B over 0.5 minutes where it was held for 1.5 minutes to re-equilibrate the column for the next sample. The data were acquired using Agilent MassHunter, version B.08.02, processed using Skyline version 4.1, and peak quantification was refined with mProphet in Skyline. All data and skyline files are available via the Panorama Public repository at this link: https://panoramaweb.org/a-platform-for-polyketide-biosynthesis-using-yeast.url. Data are also available via ProteomeXchange with identifier: PXD013388.

### Comparative metabolite profiling by LC-MS and data analysis

LC-MS analysis was performed using a Dionex Ultimate 3000 ultra-high-performance liquid chromatography (UHPLC) coupled to a UV/Vis diode array detector (DAD) and a high-resolution mass spectrometer (HRMS) Orbitrap Fusion mass spectrometer (ThermoFisher Scientific, Waltham, MA, USA). UV-Vis detection was done using a DAD-3000 in the range 200 – 700 nm. Injections of 5 µL of each sample were separated using a Zorbax Eclipse Plus C-18 column (2.1 × 100 mm, 1.8 µm) (Agilent, Santa Clara, CA, USA) at a flow rate of 0.35 mL/min, and a temperature of 35.0 °C. Mobile phases A and B were 0.1 % formic acid in water and acetonitrile, respectively. Elution was performed with a 17 min multistep system. After 5 % B for 0.3 min, a linear gradient started from 5 % B to 100 % B in 13 min, which was held for another 2 min and followed by re-equilibration to 5 % B until 17 min. HRMS was performed in ESI-, with a spray voltage of 2,750 V respectively, in the mass range range (*m/z*) 100-1,000 at a resolution of 120,000, RF Lens 50 %, and AGC target 2e5. Before analysis, the MS was calibrated using ESI Negative on Calibration Solution (P/N 88324, Thermo Scientific, San Jose, USA).

LC-MS/MS analysis was carried out using data-dependent MS/MS analysis by analyzing the most intense ions form the full-scan using a master scan time of 1.0 s. Dynamic exclusion was used to exclude ions for 20 s after two measurements within 30 s. Fragmentation was performed using stepped HCD collision energy of 15, 25, and 35% at a resolution of 30,000, RF Lens 50 %, and AGC target 1e5, while full-scan resolution was set to 60,000.

Data analyses were performed with the software Xcalibur 3.1.2412.17 (Thermo Fisher Scientific Inc.).

## Acknowledgments

We would like to acknowledge Synthetic Biology Tools for Yeast laboratory members for fruitful discussions and Anna Koza for help on sequencing efforts. This work was funded by grants from the Novo Nordisk Foundation [NNF10CC1016517], [NNF15OC0016626] and is part of the U.S. Department of Energy Joint BioEnergy Institute supported by the U.S. Department of Energy, Office of Science, Office of Biological and Environmental Research, through Contract DE-AC02-05CH11231 between Lawrence Berkeley National Laboratory and the U.S. Department of Energy.

