## Supplementary material for "Versatile polyketide biosynthesis platform for production of aromatic compounds in yeast"

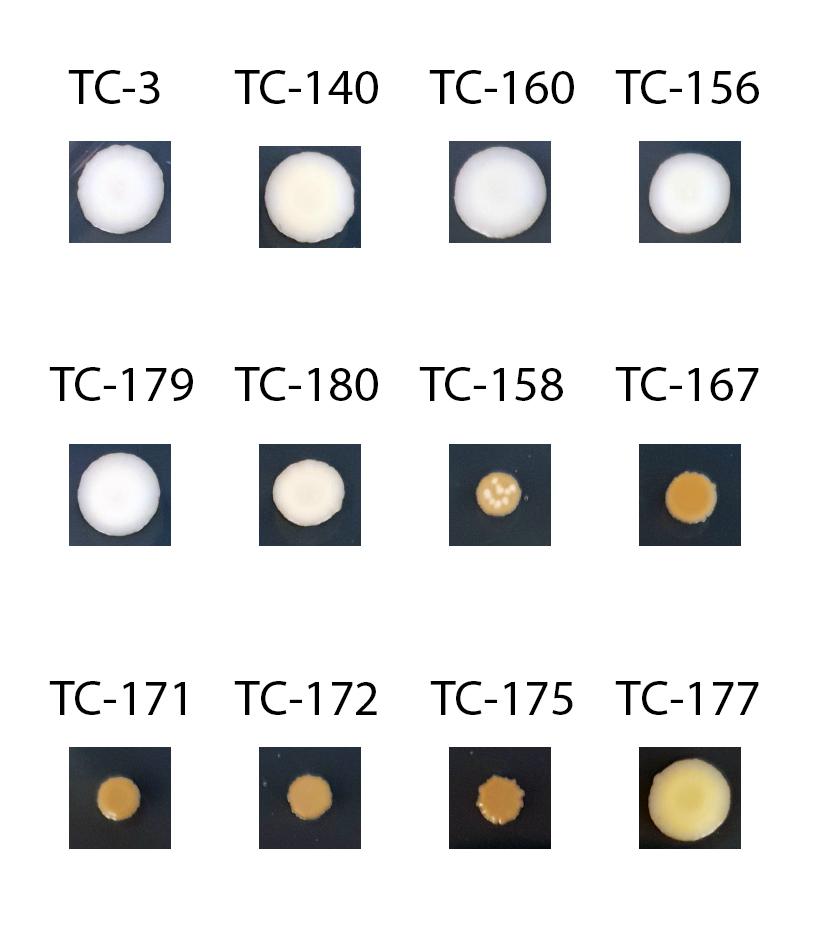

**Supplementary Figure 1**. Phenotypic comparison of constructed platform strains.

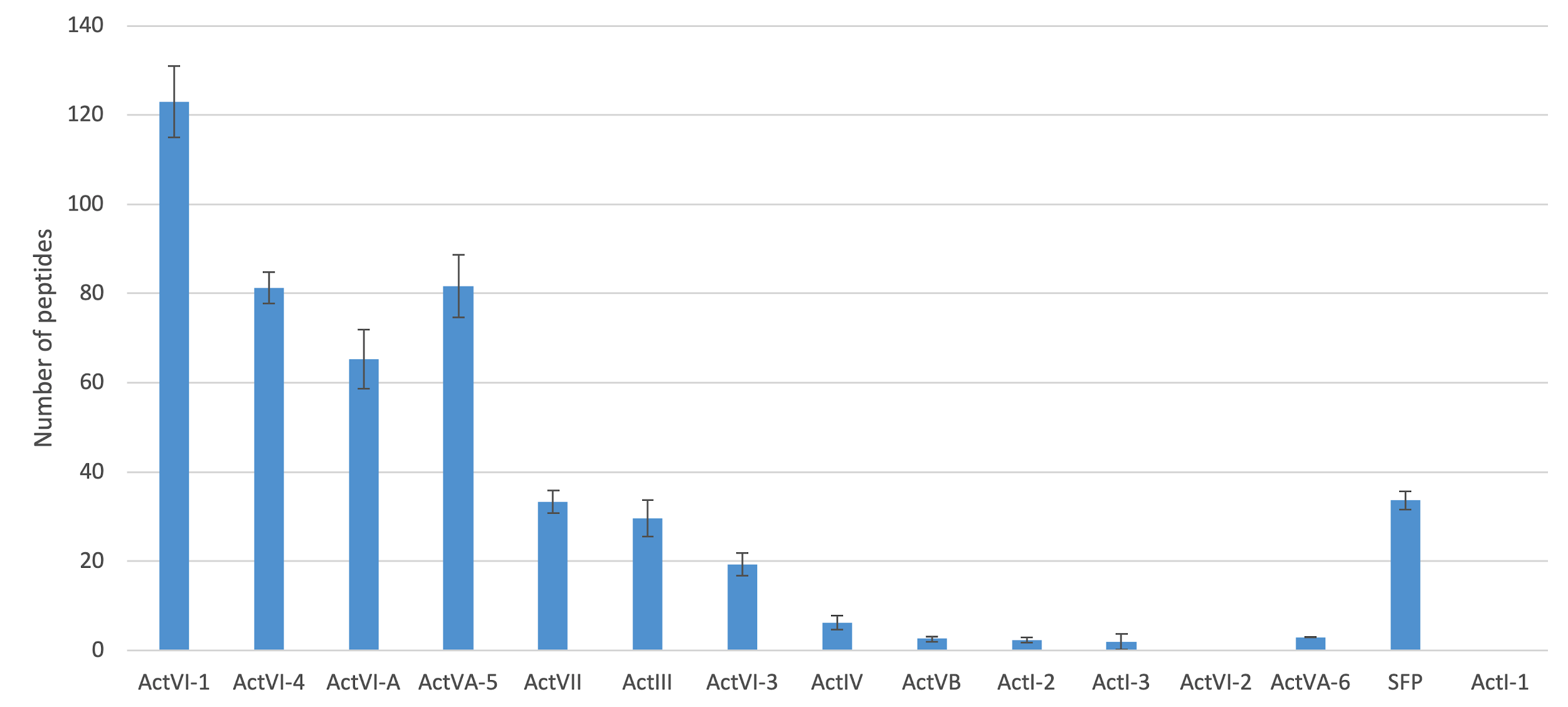
**Supplementary Figure 2**. Whole cell proteomics from strain TC-140 (expressing Act miniPKS and Act pathway). The data is based on 3 independent samples.

**a	b**
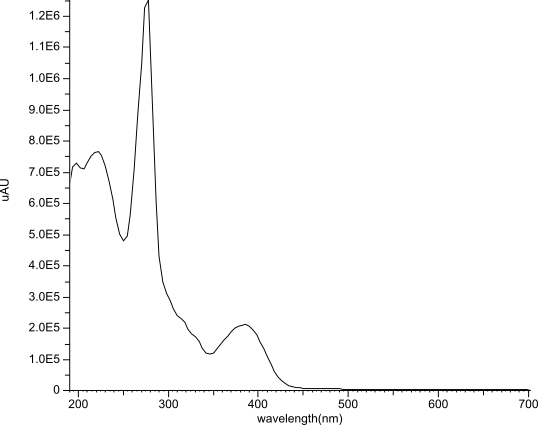

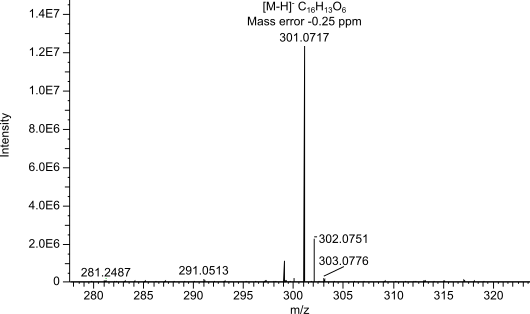

**c**

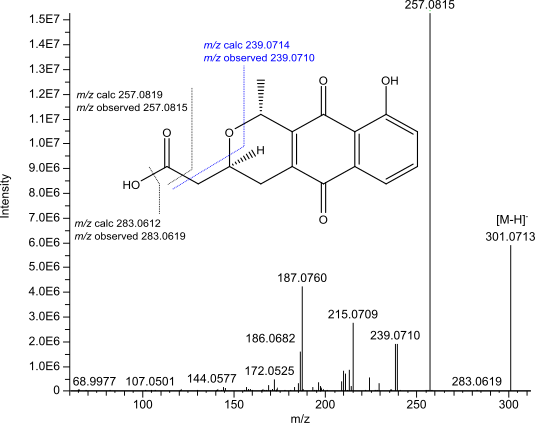

**Supplementary Figure 3**. **a**, Mass spectrum from the compound tentatively identified as the antibiotic - dihydrokalafungin (DHK). **b**, UV spectrum from the compound tentatively identified as the antibiotic - dihydrokalafungin (DHK). **c**, MS/MS spectrum from the compound tentatively identified as the antibiotic - dihydrokalafungin (DHK) with major fragments annotated. The MS/MS spectrum of DHK from yeast engineered strains matched with DHK MS/MS spectrum from *S. coelicolor.*

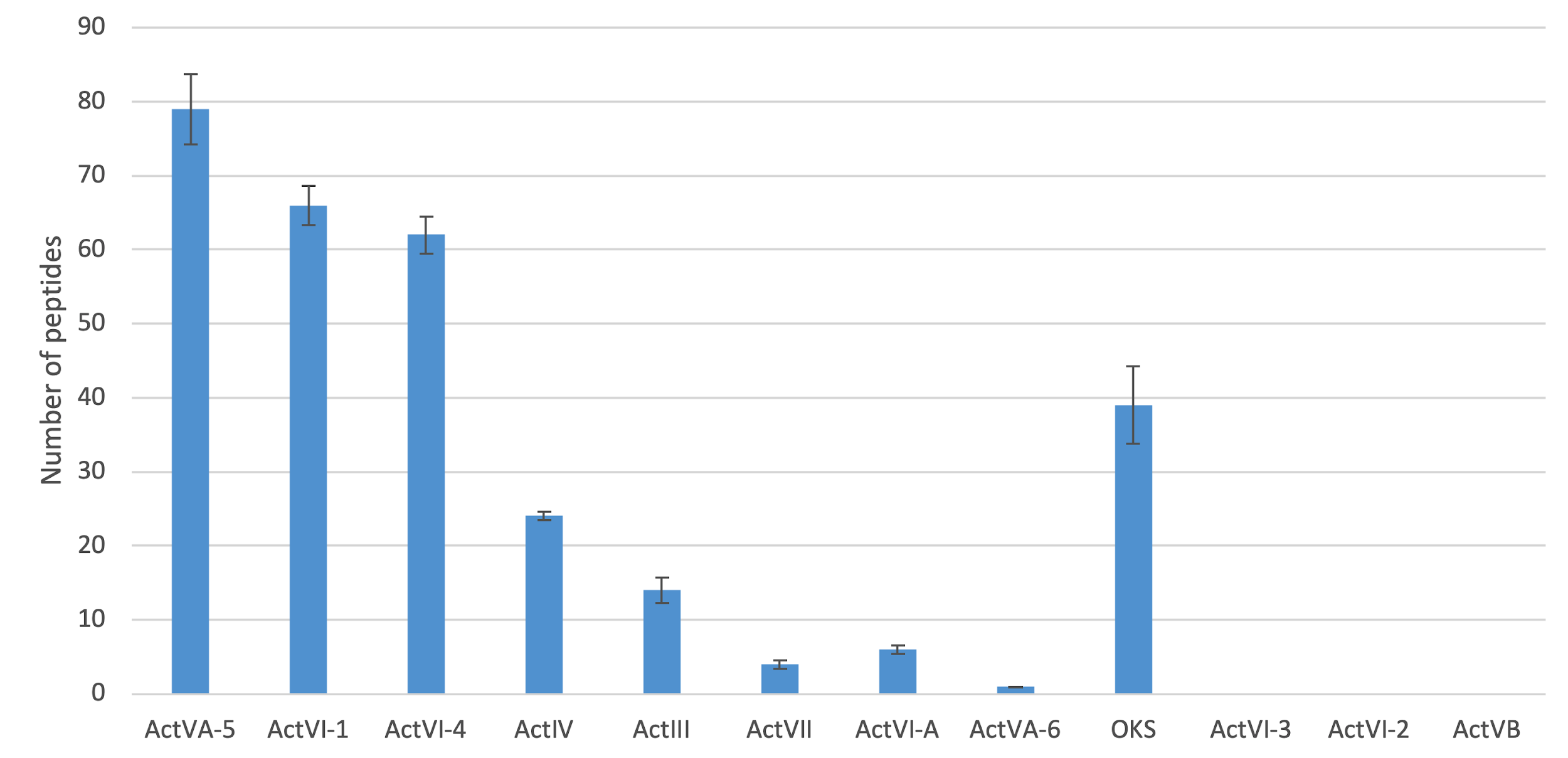

**Supplementary Figure 4**. Whole cell proteomics from strain TC-158 (expressing AaOKS and Act pathway). The data is based on 3 independent samples.

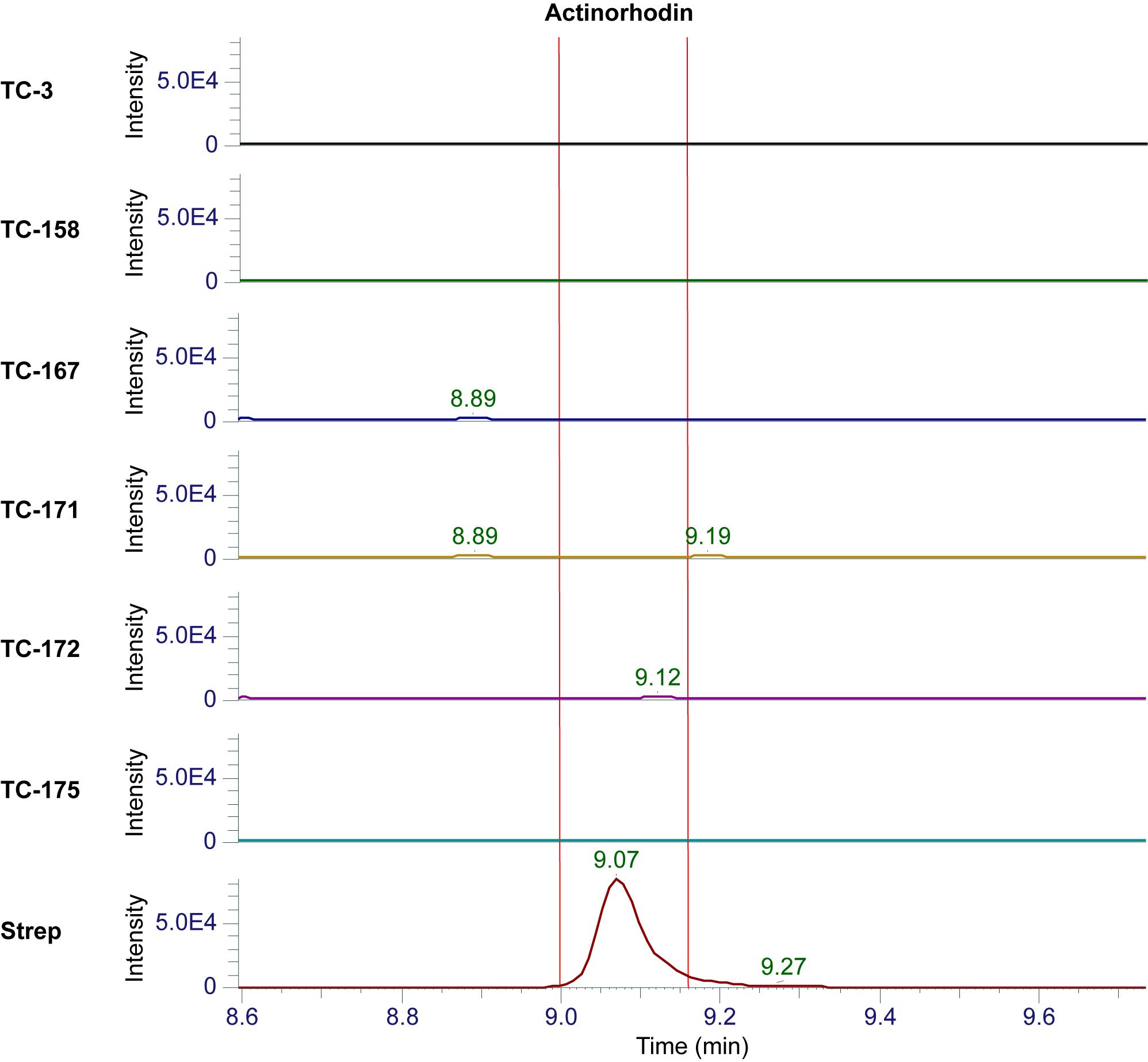

**Supplementary Figure 5**. Comparative metabolomics by LC-MS to evaluate the production of actinorhodin in engineered yeast strains.

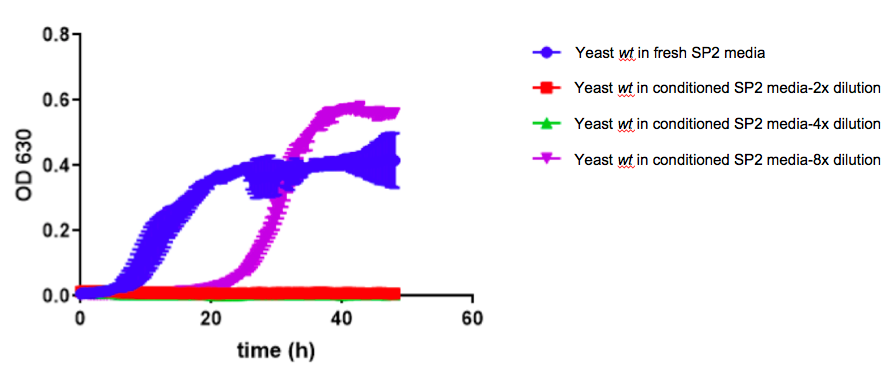

**Supplementary Figure 6**. Growth profiling of yeast *wt* strain in fresh and conditioned ISP2 media. Conditioned media was prepared by growing *Streptomyces coelicolor* in ISP2 media for 168 hours, following removal of cells by centrifugation and addition of 2% glucose. Data shown from 3 independent replicates.

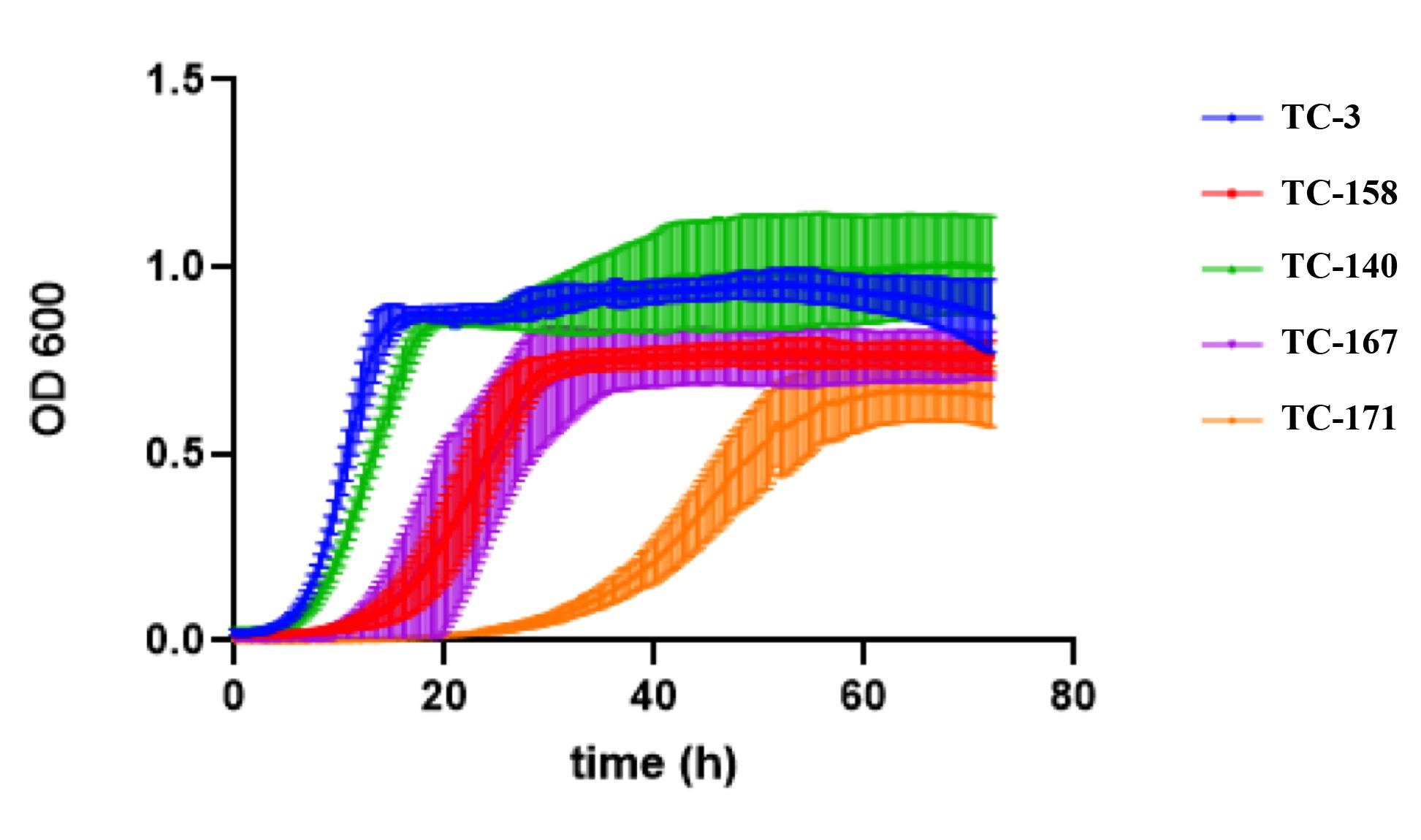

**Supplementary Figure 7**. Growth profiling of engineered yeast strains. TC-3 is *wt* control; TC-140 is strain with Act pathway expressing Act miniPKS; TC-158 strain expressing Act pathway and AaOKS; TC-167 is strain expressing AaOKS and optimized Act pathway; TC-171 is stain expressing AaOKS with Act pathway, including dimerize ActVA-4 (potential actinorhodin producer). Data shown from 3 independent replicates. Cell were grown in Synthetic Complete (SC) medium.

**Supplementary Table 1.** Construction steps, expression units and genomic integration sites for platform strains TC-140 and TC-158 respectively.

| **Step** | **Expression unit** | **Integration site** |
| --- | --- | --- |
| **Step 1** | tADH1-ActI-1 <-pTDH3-pTEF1-> ActI-3-tCYC1 | **XI-1** |
|  | tADH1-ActI-2 <-pTDH3-pTEF1-> SFP-tCYC1 | **XII-1** |
| **Step 2** | tADH1-ActIII <-pTDH3-pTEF1-> ActIV-tCYC1 | **X-3** |
|  | tADH1-ActVI-1 <-pTDH3-pTEF1-> ActVI-3-tCYC1 | **XI-2** |
|  | tADH1-ActVII <-pTDH3-pTEF1-> ActVI-A-tCYC1 | **XII-2** |
| **Step 3** | tADH1-ActVI-4 <-pTDH3-pTEF1-> ActVA-6-tCYC1 | **X-2** |
|  | pTEF1-> ActVI-2-tCYC1 | **XI-5** |
|  | tADH1-ActVA-5 <-pTDH3-pTEF1-> ActVB-tCYC1 | **XII-4** |

| **Step** | **Expression unit** | **Integration site** |
| --- | --- | --- |
| **Step 1** | pTDH3-> AaOKS-tCYC1 | **Expression from plasmid** |
| **Step 2** | tADH1-ActIII <-pTDH3-pTEF1-> ActIV-tCYC1 | **X-3** |
|  | tADH1-ActVI-1 <-pTDH3-pTEF1-> ActVI-3-tCYC1 | **XI-2** |
|  | tADH1-ActVII <-pTDH3-pTEF1-> ActVI-A-tCYC1 | **XII-2** |
| **Step 3** | tADH1-ActVI-4 <-pTDH3-pTEF1-> ActVA-6-tCYC1 | **X-2** |
|  | pTEF1-> ActVI-2-tCYC1 | **XI-5** |
|  | tADH1-ActVA-5 <-pTDH3-pTEF1-> ActVB-tCYC1 | **XII-4** |

**Supplementary Table 2**. Strain table

| Name | Expression units | Integration sites | Plasmids |
| --- | --- | --- | --- |
| TC-3 | CEN.PK2-1C + PTEF1-Cas9 |  | pRS414-TEF1p-Cas9-CYC1t |
| TC-140 | ActI-1 <-PTDH3-PTEF1-> ActI-3;  ActI-2 <-PTDH3-PTEF1-> SFP;  ActIII <-PTDH3-PTEF1-> ActIV;  ActVI-1 <-PTDH3-PTEF1-> ActVI-3;  ActVII <-PTDH3-PTEF1-> ActVI-A;  ActVI-4 <-PTDH3-PTEF1-> ActVA-6;  PTEF1-> ActVI-2;  ActVA-5 <-PTDH3-PTEF1-> ActVB. | XI-1;  XII-1;  X-3;  XI-2;  XII-2;  X-2;  XI-5;  XII-4. | pTAJAK-122;  pTAJAK-123;  pTAJAK-131;  pTAJAK-132;  pTAJAK-133;  pTAJAK-134;  pTAJAK-135;  pTAJAK-136. |
| TC-156 | ActIII <-PTDH3-PTEF1-> ActIV;  ActVI-1 <-PTDH3-PTEF1-> ActVI-3;  ActVII <-PTDH3-PTEF1-> ActVI-A;  ActVI-4 <-PTDH3-PTEF1-> ActVA-6;  PTEF1-> ActVI-2;  ActVA-5 <-PTDH3-PTEF1-> ActVB. | X-3;  XI-2;  XII-2;  X-2;  XI-5;  XII-4. | pTAJAK-131;  pTAJAK-132;  pTAJAK-133;  pTAJAK-134;  pTAJAK-135;  pTAJAK-136. |
| TC-158 | ActIII <-PTDH3-PTEF1-> ActIV;  ActVI-1 <-PTDH3-PTEF1-> ActVI-3;  ActVII <-PTDH3-PTEF1-> ActVI-A;  ActVI-4 <-PTDH3-PTEF1-> ActVA-6;  PTEF1-> ActVI-2;  ActVA-5 <-PTDH3-PTEF1-> ActVB;  PTDH3->AaOKS. | X-3;  XI-2;  XII-2;  X-2;  XI-5;  XII-4. | pTAJAK-131;  pTAJAK-132;  pTAJAK-133;  pTAJAK-134;  pTAJAK-135;  pTAJAK-136;  pTAJAK-205. |
| TC-160 | ActIII <-PTDH3-PTEF1-> ActIV;  ActVI-1 <-PTDH3-PTEF1-> ActVI-3;  ActVII <-PTDH3-PTEF1-> ActVI-A;  ActVI-4 <-PTDH3-PTEF1-> ActVA-6;  PTEF1-> ActVI-2;  ActVA-5 <-PTDH3-PTEF1-> ActVB;  PTDH3->AaOKS. | X-3;  XI-2;  XII-2;  X-2;  XI-5;  XII-4  XI-3 | pTAJAK-131;  pTAJAK-132;  pTAJAK-133;  pTAJAK-134;  pTAJAK-135;  pTAJAK-136;  pTAJAK-204. |
| TC-167 | ActIII <-PTDH3-PTEF1-> ActIV;  ActVI-1 <-PTDH3-PTEF1-> ActVI-3;  ActVII <-PTDH3-PTEF1-> ActVI-A;  ActVI-4 <-PTDH3-PTEF1-> ActVA-6;  PTEF1-> ActVI-2;  ActVA-5 <-PTDH3-PTEF1-> ActVB;  PTDH3->AaOKS;  ActVI-2 <-PTDH3-PTEF1-> ActVA-6;  ActVB <-PTDH3-PTEF1-> ActVI-3. | X-3;  XI-2;  XII-2;  X-2;  XI-5;  XII-4;  XI-1;  XII-1. | pTAJAK-131;  pTAJAK-132;  pTAJAK-133;  pTAJAK-134;  pTAJAK-135;  pTAJAK-136;  pTAJAK-205;  pTAJAK-220;  pTAJAK-221. |
| TC-171 | ActIII <-PTDH3-PTEF1-> ActIV;  ActVI-1 <-PTDH3-PTEF1-> ActVI-3;  ActVII <-PTDH3-PTEF1-> ActVI-A;  ActVI-4 <-PTDH3-PTEF1-> ActVA-6;  PTEF1-> ActVI-2;  ActVA-5 <-PTDH3-PTEF1-> ActVB;  PTDH3->AaOKS;  PTDH3->ActVA-4. | X-3;  XI-2;  XII-2;  X-2;  XI-5;  XII-4;  XII-5 | pTAJAK-131;  pTAJAK-132;  pTAJAK-133;  pTAJAK-134;  pTAJAK-135;  pTAJAK-136;  pTAJAK-205;  pTAJAK-229. |
| TC-172 | ActIII <-PTDH3-PTEF1-> ActIV;  ActVI-1 <-PTDH3-PTEF1-> ActVI-3;  ActVII <-PTDH3-PTEF1-> ActVI-A;  ActVI-4 <-PTDH3-PTEF1-> ActVA-6;  PTEF1-> ActVI-2;  ActVA-5 <-PTDH3-PTEF1-> ActVB;  PTDH3->AaOKS;  ActVI-2 <-PTDH3-PTEF1-> ActVA-6;  ActVB <-PTDH3-PTEF1-> ActVI-3;  PTDH3->ActVA-4. | X-3;  XI-2;  XII-2;  X-2;  XI-5;  XII-4;  XI-1;  XII-1;  XII-5 | pTAJAK-131;  pTAJAK-132;  pTAJAK-133;  pTAJAK-134;  pTAJAK-135;  pTAJAK-136;  pTAJAK-205;  pTAJAK-220;  pTAJAK-221;  pTAJAK-229. |
| TC-175 | ActIII <-PTDH3-PTEF1-> ActIV;  ActVI-1 <-PTDH3-PTEF1-> ActVI-3;  ActVII <-PTDH3-PTEF1-> ActVI-A;  PTDH3->AaOKS;  Med-29 <-PTDH3-PTEF1-> ActVA-6;  Med-9 <-PTDH3-PTEF1-> ActVA-4;  ActVA-5 <-PTDH3-PTEF1-> ActVB. | X-3;  XI-2;  XII-2;  X-2;  XI-5;  XII-4. | pTAJAK-131;  pTAJAK-132;  pTAJAK-133;  pTAJAK-205;  pTAJAK-227;  pTAJAK-228;  pTAJAK-136. |
| TC-177 | ActIII <-PTDH3-PTEF1-> ActIV;  ActVI-1 <-PTDH3-PTEF1-> ActVI-3;  ActVII <-PTDH3-PTEF1-> ActVI-A;  PTDH3->AaOKS;  Med-29 <-PTDH3-PTEF1-> ActVA-6  Med-9 <-PTDH3-PTEF1-> ActVA-4  Med7 <-PTDH3-PTEF1-> Med-13. | X-3;  XI-2;  XII-2;  X-2;  XI-5;  XII-4. | pTAJAK-131;  pTAJAK-132;  pTAJAK-133;  pTAJAK-205;  pTAJAK-227;  pTAJAK-228;  pTAJAK-225. |

**Supplementary Table 3**. Plasmid table

| Name | Parental plasmid | Features |
| --- | --- | --- |
| pTAJAK-122 | pCfB3036 | tADH1-ActI-1 <-pTDH3-pTEF1-> ActI-3-tCYC1 |
| pTAJAK-123 | pCfB3038 | tADH1-ActI-2 <-pTDH3-pTEF1-> SFP-tCYC1 |
| pTAJAK-131 | pCfB3034 | tADH1-ActIII <-pTDH3-pTEF1-> ActIV-tCYC1 |
| pTAJAK-132 | pCfB2903 | tADH1-ActVI-1 <-pTDH3-pTEF1-> ActVI-3-tCYC1 |
| pTAJAK-133 | pCfB3039 | tADH1-ActVII <-pTDH3-pTEF1-> ActVI-A-tCYC1 |
| pTAJAK-134 | pCfB2899 | tADH1 ActVI-4 <-pTDH3-pTEF1-> ActVA-6-tCYC1 |
| pTAJAK-135 | pCfB3037 | pTEF1-> ActVI-2-tCYC1 |
| pTAJAK-136 | pCfB3040 | tADH1-ActVA-5 <-pTDH3-pTEF1-> ActVB-tCYC1 |
| pTAJAK-204 | pCfB2904 | pTDH3-> AaOKS-tCYC1 |
| pTAJAK-205 | pESC-URA | pTDH3-> AaOKS-tCYC1 |
| pTAJAK-220 | pCfB3036 | tADH1-ActVI-2 <-pTDH3-pTEF1-> ActVA-6-tCYC1 |
| pTAJAK-221 | pCfB3038 | tADH1-ActVB <-pTDH3-pTEF1-> ActVI-3-tCYC1 |
| pTAJAK-225 | pCfB3040 | tADH1-Med-7 <-pTDH3-pTEF1-> Med-13-tCYC1 |
| pTAJAK-226 | pCfB3037 | tADH1-ActVI-2<-pTDH3-pTEF1-> ActVA-4-tCYC1 |
| pTAJAK-227 | pCfB2899 | tADH1-Med-29 <-pTDH3-pTEF1-> ActVA-6-tCYC1 |
| pTAJAK-228 | pCfB3037 | tADH1-Med-9 <-pTDH3-pTEF1-> ActVA-4-tCYC1 |
| pTAJAK-229 | pCfB2909 | pTDH3-> ActVA-4-tCYC1 |
| pTAJAK-121 | pTAJAK-71 | gRNA expression plasmid, targeting sites: XI-1, XII-1 |
| pCfB3051 | pTAJAK-71 | gRNA expression plasmid, targeting sites: X-3, XI-2, XII-2 |
| pCfB3053 | pTAJAK-71 | gRNA expression plasmid, targeting sites: X-2, XI-5, XII-4 |
| pCfB2909 | pTAJAK-71 | gRNA expression plasmid, targeting sites: XII-5 |

**Supplementary Table 4**. Primer table

| Name | Gene amplified | Sequence 5’->3’ |
| --- | --- | --- |
| TJOS-136F | ActI-1 | AGTGCAGGUATGCCACTAGATGCGGCCCC |
| TJOS-136R | ActI-1 | CGTGCGAUTCAAGCAGCAGCTCCCGCCG |
| TJOS-137F | ActI-3 | ATCTGTCAUATGGCTACATTATTGACTAC |
| TJOS-137R | ActI-3 | CACGCGAUTCAAGCAGCTTCAGCAAGTG |
| TJOS-138F | ActI-2 | AGTGCAGGUATGTCTGTTCTTATAACAGG |
| TJOS-138R | ActI-2 | CGTGCGAUTCAGGGTGTTGGTGCAAAAC |
| TJOS-139F | SFP | ATCTGTCAUATGAAGATTTACGGAATTTA |
| TJOS-139R | SFP | CACGCGAUTTATAAAAGCTCTTCGTACG |
| TJOS-140F | ActIII | AGTGCAGGUATGGCCACACAAGACAGTGA |
| TJOS-140R | ActIII | CGTGCGAUTCAGTAATTTCCTAATCCCC |
| TJOS-141F | ActIV | ATCTGTCAUATGACTGTCGAAGTCAGAGA |
| TJOS-141R | ActIV | CACGCGAUTCAAGCCAGGCAAGTTGGTA |
| TJOS-142F | ActVI-1 | AGTGCAGGUATGTCTACCGTGACAGTGAT |
| TJOS-142R | ActVI-1 | CGTGCGAUTTATTTTGTCTCTTCTTGGG |
| TJOS-143F | ActVI-3 | ATCTGTCAUATGACTTCATCATTGCATCA |
| TJOS-143R | ActVI-3 | CACGCGAUTCACTTATGTTTTTTAACCT |
| TJOS-144F | ActVII | AGTGCAGGUATGAGCAGGCCAGGTGAGCA |
| TJOS-144R | ActVII | CGTGCGAUTTAACTAGCGGGTCCGGCTG |
| TJOS-145F | ActVI-A | ATCTGTCAUATGACAATTACAGCTCTGCC |
| TJOS-145R | ActVI-A | CACGCGAUTTAGGCGGGAAAGACATGGT |
| TJOS-146F | ActVI-4 | AGTGCAGGUATGCCCAAGGCAGTCGCTAT |
| TJOS-146R | ActVI-4 | CGTGCGAUTTAAAGGTCGGGAACCAAAA |
| TJOS-147F | ActVA-6 | ATCTGTCAUATGGCTGAGGTAAATGACCC |
| TJOS-147R | ActVA-6 | CACGCGAUTTAACTAGGCAAAATAGCCC |
| TJOS-148F | ActVA-5 | AGTGCAGGUATGTCTGAAGACACTATGAC |
| TJOS-148R | ActVA-5 | CGTGCGAUTTAGCCATCGTTTGACCTCC |
| TJOS-149F | ActVB | ATCTGTCAUATGGCGGCAGATCAAGGTAT |
| TJOS-149R | ActVB | CACGCGAUCTAGCCTGCGTGCGCAGGAA |
| TJOS-150F | ActVI-2 | ATCTGTCAUATGATGAGAGCAGTCCAATT |
| TJOS-150R | ActVI-2 | CACGCGAUTTAAGGAACTAAAACAACCC |
| TJOS-368F | AaOKS | ATCTGTCAUAAAACAATGAGTAGTTTATCAAATGC |
| TJOS-368R | AaOKS | CACGCGAUTCACATCAATGGCAAGGAAT |
| TJOS-372F | ActVI-2 | AGTGCAGGUATGATGAGAGCAGTCCAATT |
| TJOS-372R | ActVI-2 | CGTGCGAUTTAAGGAACTAAAACAACCC |
| TJOS-373F | ActVB | AGTGCAGGUATGGCGGCAGATCAAGGTAT |
| TJOS-373R | ActVB | CGTGCGAUCTAGCCTGCGTGCGCAGGAA |
| TJOS-375F | ActVA-4 | ATCTGTCAUATGCCGGATGAAAATAAGCC |
| TJOS-375R | ActVA-4 | CACGCGAUTTATGCAGGGTCCCTGGGAG |
| TJOS-376F | Med-7 | AGTGCAGGUATGCCTGCCACTCAGCCAAC |
| TJOS-376R | Med-7 | CGTGCGAUTTAAGGAGTATCCAAGGCTC |
| TJOS-377F | Med-29 | AGTGCAGGUATGTTATCTGTCGTGATGGA |
| TJOS-377R | Med-29 | CGTGCGAUTTATCTGTCTCCGTTGGTCG |
| TJOS-378F | Med-9 | AGTGCAGGUATGCCTTGTGCATTGTCTGA |
| TJOS-378R | Med-9 | CGTGCGAUTTAGGCGCCGCCTCCTGCAG |
| TJOS-381F | Med-13 | ATCTGTCAUATGACCGTGAGAGCCGACAT |
| TJOS-381R | Med-13 | CACGCGAUTTACGTCGCAGTACGGAACC |
